# Antigen presentation requirements for effective cDC1-based cancer immunotherapy

**DOI:** 10.64898/2026.01.24.701412

**Authors:** Josué E. Pineda, Tomoyuki Minowa, Li Shen, Yifan Zhou, Allison M. Dyevoich, Bhakti Patel, Sarah M. Schneider, Sunita Keshari, Akata Saha, Morgan N. Riba, Jing Wang, Stephanie S. Watowich, Matthew M. Gubin

## Abstract

Type 1 conventional dendritic cells (cDC1s) are important for generating and sustaining antitumor immunity. Accordingly, the abundance of cDC1s in human tumors correlates with improved outcomes in cancer. Capitalizing on this role, we previously demonstrated that vaccination with *in vitro*-derived murine cDC1s elicits durable tumor control in multiple preclinical models; however, the immunological mechanisms underlying the efficacy of cDC1 vaccination remain unclear. Here, we examined whether *in vitro*-derived cDC1s resemble tumor-infiltrating DC populations and whether MHC-I and MHC-II antigen presentation contribute to cDC1-mediated tumor control following vaccination in melanoma. As expected, MHC-I- or MHC-II-deficiency had minimal impact on the transcriptional state of cDC1s in homeostasis or following stimulation with the adjuvant poly dI:dC. Moreover, *in vitro*-derived cDC1s cultured under steady-state conditions closely resembled tumor-infiltrating cDC1s, whereas their poly dI:dC-stimulated counterparts resembled CCR7^+^ tumor-infiltrating DC populations, also referred to as mregDCs or LAMP3^+^ DCs. Our data further show that both MHC-I and MHC-II contribute to tumor control upon cDC1 vaccination, and coexpression of MHC-I and MHC-II on the same cDC1 is necessary for a robust vaccine response. We also identified an important function for host cDC1s in supporting the efficacy of vaccination with *in vitro*-derived cDC1s, as judged by impaired tumor control in *Irf8+32**^-/-^*** mice, which lack endogenous cDC1s. Overall, these results indicate that effective antitumor responses depend on MHC-I and MHC-II antigen presentation by vaccine-delivered cDC1s, with additional contributions from host cDC1s.

**Key points:**

- *In vitro*-generated cDC1s resemble intratumoral DC populations found in mice and humans.
- MHC-I and MHC-II antigen presentation by vaccine-delivered cDC1s contribute to antitumor efficacy.
- Coexpression of MHC-I and MHC-II on the same cDC1 enhances vaccine responses.
- Antitumor responses reflect the activity of vaccine and endogenous cDC1s.

## Introduction

Dendritic cells (DCs) have gained attention in cancer immunotherapy research because of their exceptional capacity to present tumor antigens and coordinate T cell-mediated immunity [1–6]. Several clinical strategies have explored DC-based vaccines, including those using monocyte-derived DCs (MoDC) and *in vivo* antigen-targeting approaches that exploit DC-specific receptors [4, 7–11]. One of the earliest personalized neoantigen vaccine clinical trials utilized MoDCs, finding that these vaccines could elicit *de novo* tumor antigen-specific T cell responses and boost pre-existing responses in patients with melanoma [7]. However, despite measurable immunogenicity across these different approaches, DC-based cancer vaccines have achieved limited clinical efficacy, emphasizing the need to improve DC-based therapeutic strategies [12, 13].

Among DC populations, type 1 conventional DCs (cDC1s) are a specialized subset with optimal ability to cross-present exogenous antigens on MHC class I (MHC-I) to prime CD8^+^ T cells, a process essential for tumor rejection [6, 14–16]. The development of cDC1s depends on the transcription factors IRF8, ID2, and BATF3, as well as signaling through the Fms-like tyrosine kinase 3 receptor (FLT3) in response to its ligand Flt3L, which are critical for cDC1 differentiation *in vivo* and *in vitro* [15]. Under homeostatic conditions, cDC1s reside within lymphoid organs and peripheral tissues such as the skin, lung, and intestine, where they continuously sample and present antigens. Tissue-resident cDC1s are capable of undergoing homeostatic or pathogen-induced migration to lymph nodes (LNs), enabling cDC1s to regulate tolerogenic or adaptive immune responses via antigen presentation to T cells [15].

cDC1s act as central coordinators of antitumor immunity through multiple complementary mechanisms. In tumor-draining LNs (TdLNs), cDC1s provide canonical antigen presentation and costimulatory signals to activate naïve antitumor T cell populations. cDC1s are also proficient in cross-presentation of exogenous tumor antigens on MHC-I to stimulate CD8^+^ T cell responses. Simultaneously, cDC1s present antigen on MHC-II to engage CD4^+^ T cells. Prior studies show that upon CD4^+^ T cell engagement, CD4^+^ T cells license cDC1s through CD40 ligand (CD40L)-CD40 interactions, which enhances the cDC1’s ability to prime cytotoxic CD8^+^ T cells and sustain productive antitumor immunity [17]. In addition to their antigen-presenting functions, cDC1s secrete cytokines such as interleukin-12 (IL-12) and type I interferons that promote T cell activation, survival, and effector differentiation, driving polarization of IFN-γ^+^ CD4^+^ T cells (Th1 cells) and CD8^+^ T cell cytotoxic responses [18–20]. Within the tumor microenvironment (TME), cDC1s facilitate the recruitment and localization of effector T cells through the production of chemokines such as CXCL9 and CXCL10 [20–22]. Accordingly, the abundance and activation of cDC1s within tumors correlates with enhanced antitumor immune activity, increased responsiveness to immune checkpoint blockade, and improved clinical outcomes across multiple cancer types [1, 3, 4, 15, 16, 23–27]. However, many tumors exhibit a profound deficiency or dysfunction in cDC1s, contributing to limited T cell infiltration and inadequate immune control [25, 28]. Therefore, strategies that enhance cDC1 activation, antigen presentation, and persistence within the TME hold considerable potential to improve cancer immunotherapy outcomes.

We previously established a cDC1-based vaccine using cells generated *in vitro* from murine bone marrow [23, 25, 26, 29]. Our studies demonstrated that delivery of adjuvant-activated, antigen-loaded cDC1s to the TME elicits systemic and durable antitumor immunity; moreover, intravenous delivery of the cDC1 vaccine restrains outgrowth of distal metastases. We also showed that cDC1 vaccination has superior tumor control relative to MoDC vaccination and demonstrated efficacy against a variety of murine solid tumors, suggesting tumor-agnostic activity [23, 25, 26, 29]. Recent reports from others have confirmed the efficacy of cDC1 vaccination and demonstrated its superiority to vaccination with type 2 cDCs (cDC2s) [30]. Despite these advances, the immunological mechanisms by which cDC1 vaccination elicits robust tumor control remain unknown. Here, we investigate whether MHC-I and MHC-II antigen presentation and the presence of endogenous cDC1s contribute to the efficacy of adjuvant-activated, tumor antigen-loaded cDC1 vaccines.

## Materials and methods

### Mouse strains

All mice used were obtained from The Jackson Laboratory (JAX, Bar Harbor, ME, USA) and were on the C57BL/6 background. These include C57BL/6J wild-type (WT; JAX Stock No. 000664), B6.129P2-*H2-K1^b-tm1Bpe^ H2-D1^b-tm1Bpe^*/DcrJ (MHC-I^KO^; JAX Stock No. 019995), B6.129S2-*H2^dlAb1-Ea^*/J (MHC-II^KO^; JAX Stock No. 003584), C57BL/6-*Rr172^em1Kmm^*/J (*Irf8+32*^-/-^; JAX Stock No. 032744), and C57BL/6-Tg(TcraTcrb)1100Mjb/J (OT-I; JAX Stock No. 003831) mice [31–36]. To generate MHC-I^KO^ mice, B6.129P2-*H2-K1^b-tm1Bpe^ H2-D1^b-tm1Bpe^*/DcrJ sperm was retrieved from cryopreservation and implanted into C57BL/6J mice by JAX. Progeny were obtained and bred in-house to generate double-knockout mice lacking *H2*-*K1* and *H2*-*D1* (H2-K^b^ and H2-D^b^). All experiments were conducted using 8- to 12-week-old mice. Mice were maintained under specific pathogen-free conditions at The University of Texas MD Anderson Cancer Center (Houston, TX) and used in accordance with Institutional Animal Care and Use Committee (IACUC) approved protocols.

### *In vitro* generation of cDC1s

Culture conditions to generate cDC1s were adapted from a protocol developed by Mayer *et al*. [37] and performed using methods previously described by our laboratory [23, 25, 26, 29]. Murine bone marrow cells (1.5 x 10^6^ cells/mL) were cultured in RPMI 1640 containing 10% heat-inactivated fetal bovine serum (FBS; Atlanta Biologics, Atlanta, GA, USA), 1% penicillin-streptomycin, 1 mM sodium pyruvate, and 50 µM β-mercaptoethanol (complete RPMI), and supplemented with 2 ng/mL murine granulocyte-macrophage colony-stimulating factor (mGM-CSF; PeproTech, Rocky Hill, NJ, USA) and 50 ng/mL human Flt3L (hFlt3L; PeproTech). On day 9, non-adherent cells were collected and replated in fresh complete RPMI at 0.3 x 10^6^ cells/mL with 2 ng/mL mGM-CSF and 50 ng/mL hFlt3L. On days 15-17, non-adherent cells were collected and cDC1s (CD11c^+^ CD45R^-^ CD24^+^ CD172α^-^ CD103^+^) were purified by fluorescence-activated cell sorting (FACS) using a FACSAria or FACSAria Fusion cell sorter (BD Biosciences, Palo Alto, CA, USA).

### Preparation and administration of cDC1 vaccines

FACS-purified cDC1s were cultured (2-4.5 x 10^6^ cells/mL) in complete RPMI supplemented with 20 ng/mL mGM-CSF, 20 µg/mL poly(deoxyinosinic-deoxycytidylic) acid (poly dI:dC; Sigma-Aldrich, St. Louis, MO, USA), and 200 µg/mL ovalbumin (OVA; Sigma-Aldrich) for 4 hours at 37°C. Following stimulation, cells were washed three times with phosphate-buffered saline (PBS) and resuspended in endotoxin-free PBS. For intratumoral vaccination, 3 x 10^6^ cDC1s were injected directly into established tumors of mice bearing OVA-expressing B16-F10 melanoma (B16-OVA) on days 4 and 7 after tumor implantation, as previously described [26]. For mixed vaccines (Mix^KO^), mice received a total of 6 x 10^6^ cDC1s per treatment, composed of equal numbers of MHC-I^KO^ cDC1s and MHC-II^KO^ cDC1s (3 x 10^6^ each). For experiments assessing CD40 stimulation, cDC1s were stimulated with 20 ng/mL mGM-CSF, 20 µg/mL poly dI:dC, and 200 µg/mL OVA, in the presence or absence of 20 µg/mL agonistic anti-CD40 antibody (αCD40; Leinco Technologies, Fenton, MO, USA). Cells were washed with PBS and prepared for intratumoral injection as described above.

### Tumor transplantation and monitoring

Murine B16-OVA melanoma cells, originally generated by transfection of B16-F10 cells with a plasmid encoding full-length OVA under a constitutive promoter [38], were maintained in Dulbecco’s modified Eagle medium (DMEM; Gibco, Grand Island, NY, USA) supplemented with 10% FBS and 1% penicillin-streptomycin (complete DMEM). Cells were passaged 3-5 times prior to use. WT C57BL/6J, MHC-I^KO^, MHC-II^KO^, or *Irf8+32*^-/-^ mice were shaved and injected subcutaneously on the flank with 3 x 10^5^ B16-OVA cells in 150 µL of endotoxin-free PBS. Tumor length and width were measured every 2-3 days using digital calipers. Mice were euthanized when tumor diameters reached 15 mm in any direction, or when ulceration occurred, in accordance with IACUC-approved protocols.

### *In vitro* activation and phenotypic analysis of cDC1s

FACS-purified cDC1s were cultured at 2-4.5 x 10^6^ cells/mL in complete RPMI supplemented with 20 µg/mL poly dI:dC, 20 ng/mL mGM-CSF, and 200 µg/mL OVA ± 20 µg/mL αCD40 for 4, 16, or 24 hours at 37°C, or were left untreated. Following culture, cells were washed three times with PBS and incubated with Fc block (anti-mouse CD16/32; clone 2.4G2, Tonbo Biosciences, San Diego, California, USA) in FACS buffer (PBS supplemented with 2% FBS and 2 mM ethylenediaminetetraacetic acid (EDTA)) for 15 minutes at 4°C. Cells were then stained with master mixes comprising fluorescent antibodies specific for murine surface markers for 15-30 minutes at 4°C, washed in FACS buffer, and prepared for flow cytometric analysis. Antibodies used for phenotypic analysis included APC-conjugated CD70 (FR70) and CD45R/B220 (RA3-6B2); BV605-conjugated CD103 (2E7); BV711-conjugated CD40 (3/23), CD24 (M1/69), and MHC-II (I-A/I-E; M5/114.15.2); BV785-conjugated CD86 (GL-1); eFluor^TM^ 450-conjugated CD11c (N418); FITC-conjugated CD172α/Sirpα (P84) and H-2K^b^ (AF6-88.5); PE-Cy7-conjugated CD70 (FR70) and CD80 (16-10A1); PerCP-Cy5.5-conjugated CD40 (3/23) and CD24 (M1/69); and BV650-conjugated MHC-II (I-A/I-E; M5/114.15.2). All antibodies were purchased from Tonbo Biosciences, BD Biosciences, Thermo Fisher Scientific (Waltham, MA, USA), or BioLegend (San Diego, CA, USA). Dead cells were excluded using Ghost Dye^TM^ Violet 510 (Tonbo Biosciences). Samples were acquired on a BD LSRFortessa flow cytometer (BD Biosciences), and data were analyzed using FlowJo v10 software (FlowJo, Ashland, OR, USA).

### Antigen presentation assays

FACS-purified cDC1s were stimulated with 20 µg/mL poly dI:dC and 20 ng/mL mGM-CSF, and pulsed with either 200 µg/mL OVA or 200 µg/mL of OVA-I-SIINFEKL peptide (Peptide 2.0, Chantilly, VA, USA, custom order) for 2 hours at 37°C. Cells were then washed three times with PBS. OVA-specific CD8^+^ T cells were isolated by FACS from single cell suspensions of spleens from OT-I mice and labeled with 5-(and-6)-carboxyfluorescein diacetate succinimidyl ester (CFSE; Tonbo Biosciences, San Diego, CA, USA) according to the manufacturer’s instructions for 10-20 minutes at room temperature in the dark. Labeled OT-I CD8^+^ T cells were washed twice with endotoxin-free PBS and co-cultured with poly dI:dC-stimulated and antigen-loaded cDC1s for 48 or 60 hours at 37°C in complete RPMI. Following co-culture, cells were washed three times with PBS, incubated with Fc block for 15-30 minutes at 4°C, and stained with fluorescently conjugated antibodies for murine surface markers for 15-30 minutes at 4°C. Antibodies used included APC-conjugated CD45R/B220 (RA3-6B2) and CD8α (53-6.7); BV605-conjugated CD103 (2E7); BV711-conjugated CD24 (M1/69); eFluor^TM^ 450-conjugated CD11c (N418); FITC-conjugated CD172α/Sirpα (P84); PerCP-Cy5.5-conjugated CD24 (M1/69); BV785-conjugated CD3 (17A2); PE-594-conjugated CD152 (UC10-4B9); and PE-conjugated CD279 (29F.1A12). All antibodies were purchased from Tonbo Biosciences, BD Biosciences, Thermo Fisher Scientific, or BioLegend. Dead cells were excluded with Ghost Dye^TM^ Violet 510. OT-I splenocytes and cDC1s were sorted using a FACSAria or FACSAria Fusion. Samples were acquired on a BD LSRFortessa flow cytometer and analyzed using FlowJo v10 software.

### RNA isolation, library preparation, and sequencing

Total RNA was isolated from FACS-purified cDC1s using the Qiagen RNeasy Mini Kit (Qiagen, Hilden, Germany) according to the manufacturer’s instructions. Purified RNA was stored at -80°C until shipment for sequencing. Individual biological samples were submitted to Novogene Corporation Inc. (Sacramento, CA, USA) for quality assessment, library preparation, and sequencing. RNA concentration and purity were evaluated by NanoDrop spectrophotometry, 1% agarose gel electrophoresis, and RNA integrity was assessed using an Agilent 2100 Bioanalyzer to determine RNA Integrity Numbers (RIN), a standardized quantitative measure of RNA degradation based on electrophoresis profiles [39]. Libraries were generated using a poly(A)-enriched mRNA workflow and sequenced on an Illumina NovaSeq X-Plus platform to produce 150-bp paired-end reads.

### Bioinformatics processing and differential expression analysis

For data processing, raw sequencing reads were processed using the nf-core/rnaseq pipeline v3.14 [40], a community-curated workflow built upon the nf-core framework [41]. The workflow was executed with Nextflow v23.10.1 [42] under reproducible software environments provided through Bioconda [43] and BioContainers [44]. Reads were aligned to the GRCm39 reference genome using STAR, and transcript quantification was performed with RSEM to generate gene-level count estimates. Default nf-core parameters were used unless otherwise specified. Differentially expressed genes (DEGs) were identified using the DESeq2 package, which models count data with a negative binomial framework. DESeq2 estimates gene-wise dispersions and log2 fold changes using empirical priors to improve stability, particularly for low-count genes. Statistical significance was assessed using Wald tests and resulting p-values were adjusted for multiple testing using the Benjamini-Hochberg procedure to control the false discovery rate (FDR).

### Projection of bulk RNA-seq data to scRNA-seq

Processed single-cell RNA-sequencing (scRNA-seq) datasets of tumor-infiltrating human and mouse DCs were used as reference datasets for projection analyses [45]. Projection from bulk RNA-sequencing (bulk RNA-seq) results from *in vitro*-derived cDC1s onto this reference was performed using the projectLSI pipeline (version 0.1.0) [46]. Bulk RNA-seq data were first converted into pseudo-single-cell data using the CreateSeuratObject function and normalized with the NormalizeData function in Seurat (version 5.1.0). For projection onto the human scRNA-seq reference, mouse gene symbols were converted to their human orthologs using the convert_mouse_to_human_symbols function in nichenetr (version 2.1.5). Both the scRNA-seq reference and the pseudo-single-cell dataset were processed by computing latent semantic indexing (LSI) embeddings using the calcLSI function, followed by generation of a low-dimensional representation using the CreateDimReducObject function within the projectLSI workflow. UMAP visualization was performed using the umap function (n_neighbors = 20) implemented in uwot (version 0.2.2). Then, the reference scRNA-seq and pseudo-single-cell datasets were merged, and the joint UMAP embedding was used for visualization. DEGs in the reference scRNA-seq dataset were identified using the FindAllMarkers function in Seurat and visualized with DoHeatmap, enabling annotation of DC subsets including cDC1, cDC2, and CCR7⁺ DCs.

### Statistical analyses

All statistical analyses were performed using GraphPad Prism version 10 (GraphPad Software, San Diego, CA, USA). The statistical tests applied to each dataset, including multiple and survival analyses, are specified in the corresponding figure legends. Results were considered significant when p < 0.05.

## Results

### Transcriptional similarities between *in vitro*-derived cDC1s and intratumoral DCs

To test the importance of MHC-I and MHC-II expression in vaccine-delivered cDC1s, we used a previously established culture system to generate cDC1s from the bone marrow of wild-type (WT) C57BL/6J mice, H2-K^b-/-^ H2-D^b-/-^ (MHC-I^KO^) mice, and B6.129S2-*H2^dlAb1-Ea^*/J (MHC-II^KO^) mice (**Figure 1A**) [31–35, 37]. Flow cytometric analysis confirmed the expected loss of MHC-I or MHC-II surface expression on MHC-I^KO^ or MHC-II^KO^ cDC1s, respectively (**Figure 1B**). Comparable numbers of cDC1s were derived from mice of all genotypes, with high purity, and stimulation with poly dI:dC enhanced surface expression of CD80 and CD86 similarly between WT and MHC-I^KO^ or WT and MHC-II^KO^ cDC1s (**Figures S1A-C**). In addition, we did not detect differences in the expression of MHC-I or MHC-II on WT and MHC-II^KO^ or WT and MHC-I^KO^ cDC1s, respectively (**Figure S1C**). These data indicate that loss of MHC-I or MHC-II does not impair cDC1 differentiation, expansion, or poly dI:dC-induced maturation *in vitro*.

**Figure 1.**
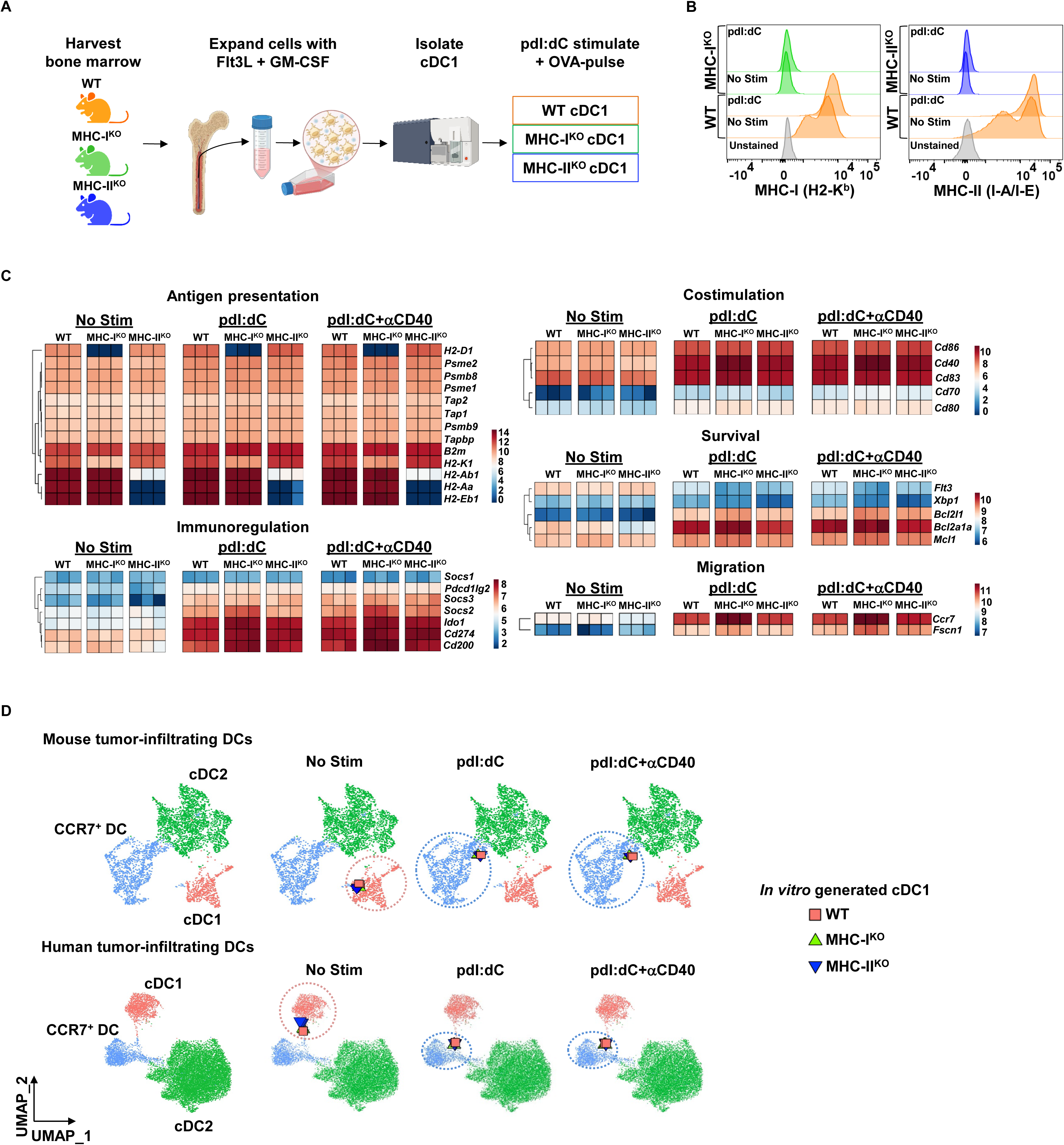
Characterization and functional evaluation of WT, MHC-I^KO^, MHC-II^KO^ cDC1s. (**A**) Schematic of the workflow used to generate cDC1s from WT, MHC-I^KO^, or MHC-II^KO^ mice *in vitro*. BM cells were cultured in medium containing Flt3L and GM-CSF, followed by FACS purification, poly dI:dC stimulation, and incubation with OVA. (**B**) Flow cytometry histogram plots showing cell surface expression of MHC-I (H2-K^b^) and MHC-II (I-A/I-E) on cDC1s after 16 hours under the indicated conditions. n = 6 (WT), n = 2 (MHC-I^KO^), n = 4 (MHC-II^KO^); representative data from at least two independent experiments. (**C**) Bulk RNA-seq analysis of cDC1s in steady-state conditions (no stim) or following stimulation with poly dI:dC with or without αCD40, as indicated. The heatmaps show log_2_(TPM + 1) gene expression, grouped according to cDC1 functions. (**D**) Mapping of bulk RNA-seq profiles from *in vitro*-derived cDC1s onto mouse or human tumor-infiltrating DC scRNA-seq reference datasets using ProjectLSI. Reference scRNA-seq cells are displayed as dots, and bulk RNA-seq samples are overlaid as shapes, corresponding to individual genotypes, as indicated.

To evaluate whether MHC-I- or MHC-II-deficiency intrinsically alters the transcriptional identity or activation of cDC1s, we performed RNA-seq using *in vitro*-derived WT, MHC-I^KO^, and MHC-II^KO^ cDC1s cultured under steady-state conditions or following stimulation with poly dI:dC, either alone or in combination with agonistic CD40 antibody (αCD40). As expected, MHC-I^KO^ cDC1s displayed defects in *H2-D1* and *H2-K1* transcript expression, while MHC-II^KO^ cDC1s showed loss of major MHC-II genes (**Figure 1C**). In contrast, WT, MHC-I^KO^, and MHC-II^KO^ cDC1s expressed comparable levels of genes involved in antigen processing and presentation, including proteasome activators (*Psme1* and *Psme2*), TAP transporters (*Tap1* and *Tap2*), and tapasin (*Tapbp*) (**Figure 1C**), suggesting that protein degradation and peptide-loading pathways relevant to antigen presentation remained intact. Stimulation with poly dI:dC ± αCD40 induced similar transcriptional responses among WT, MHC-I^KO^, and MHC-II^KO^ cDC1s, with no major differences detected in genes associated with costimulation, survival, or migration (**Figure 1C**).

Examination of the top 20 most variable genes across all conditions further indicated that transcriptional patterns were broadly similar between WT, MHC-I^KO^, and MHC-II^KO^ cDC1s **(Figure S1D**). A notable exception was *Btnl2*, which encodes butyrophilin-like 2 (BTNL2); *Btnl2* was upregulated in MHC-II^KO^ cDC1s relative to WT or MHC-I^KO^ cDC1s. Given prior reports implicating BTNL2 in the dampening of T cell activation and proliferation [47, 48], we tested the ability of MHC-II^KO^ cDC1s to induce CD8^+^ T cell proliferation. In co-culture assays with OT-I CD8^+^ T cells, MHC-II^KO^ cDC1s induced CD8^+^ T cell proliferation to a similar extent as WT cDC1s, suggesting that *Btnl2* upregulation does not impair their ability to promote CD8^+^ T cell division (**Figure S1E**). Consistent with this observation, the proportion of PD-1^+^ OT-I CD8^+^ T cells, and the magnitude of PD-1 expression were comparable between CD8^+^ T cells from co-cultures with WT or MHC-II^KO^ cDC1s (**Figures S1F-G**). Together, these data indicate that WT and MHC-II^KO^ cDC1s are similarly capable of inducing CD8^+^ T cell activation and proliferation *in vitro*.

To assess whether *in vitro*-derived cDC1s resemble naturally occurring DC populations, we compared their transcriptional profiles to human and mouse scRNA-seq reference datasets of tumor-infiltrating DCs derived from multiple cancer types and independent studies (**Figure 1D**) [45]. Bulk RNA-seq profiles from *in vitro*-derived cDC1s were embedded together with scRNA-seq data into shared UMAP space, revealing close alignment with *bona fide* intratumoral DC populations. Canonical cDC1, cDC2, and CCR7⁺ DC subsets within the UMAP were annotated based on their DEG signatures (**Figure S1H**). In this analysis, *in vitro*-derived cDC1s in homeostatic conditions were transcriptionally similar to tumor-infiltrating cDC1s, whereas cDC1s stimulated with poly dI:dC, with or without αCD40, aligned with intratumoral CCR7⁺ DCs. The CCR7⁺ DCs are also referred to as mature DCs enriched in immunoregulatory molecules (mregDCs) or lysosomal-associated membrane protein 3 (LAMP3) DCs [49–51]. Collectively, these data demonstrate that murine cDC1s generated *in vitro* from bone marrow cultures closely recapitulate transcriptional states observed in tumor-infiltrating DC populations in both mice and humans, and that loss of MHC-I or MHC-II does not substantially alter cDC1 identity, maturation, or transcriptional activation *in vitro*.

### MHC-I- and MHC-II-dependent antigen presentation contribute to the therapeutic efficacy of cDC1 vaccines

We next tested whether MHC-I or MHC-II expression on vaccine-delivered cDC1s is required for tumor control following cDC1 vaccination. WT mice were implanted with B16-OVA melanoma cells; OVA contains both MHC-I and MHC-II-restricted antigens. Using our established vaccination approach [26], mice were then treated intratumorally on days 4 and 7 with poly dI:dC-activated, OVA-loaded WT, MHC-I^KO^, or MHC-II^KO^ cDC1s or with PBS (**Figure 2A**). Consistent with our prior studies [26], vaccination with WT cDC1s significantly restrained tumor growth and prolonged mouse survival compared to PBS-treated controls (**Figures 2B-2D**). By contrast, vaccination with MHC-I^KO^ or MHC-II^KO^ cDC1s modestly controlled tumor growth, relative to the PBS treatment group, yet this was suboptimal relative to WT cDC1s. These data indicate that effective tumor control by cDC1 vaccination requires MHC-I and MHC-II antigen presentation by vaccine-delivered cDC1s.

**Figure 2.**
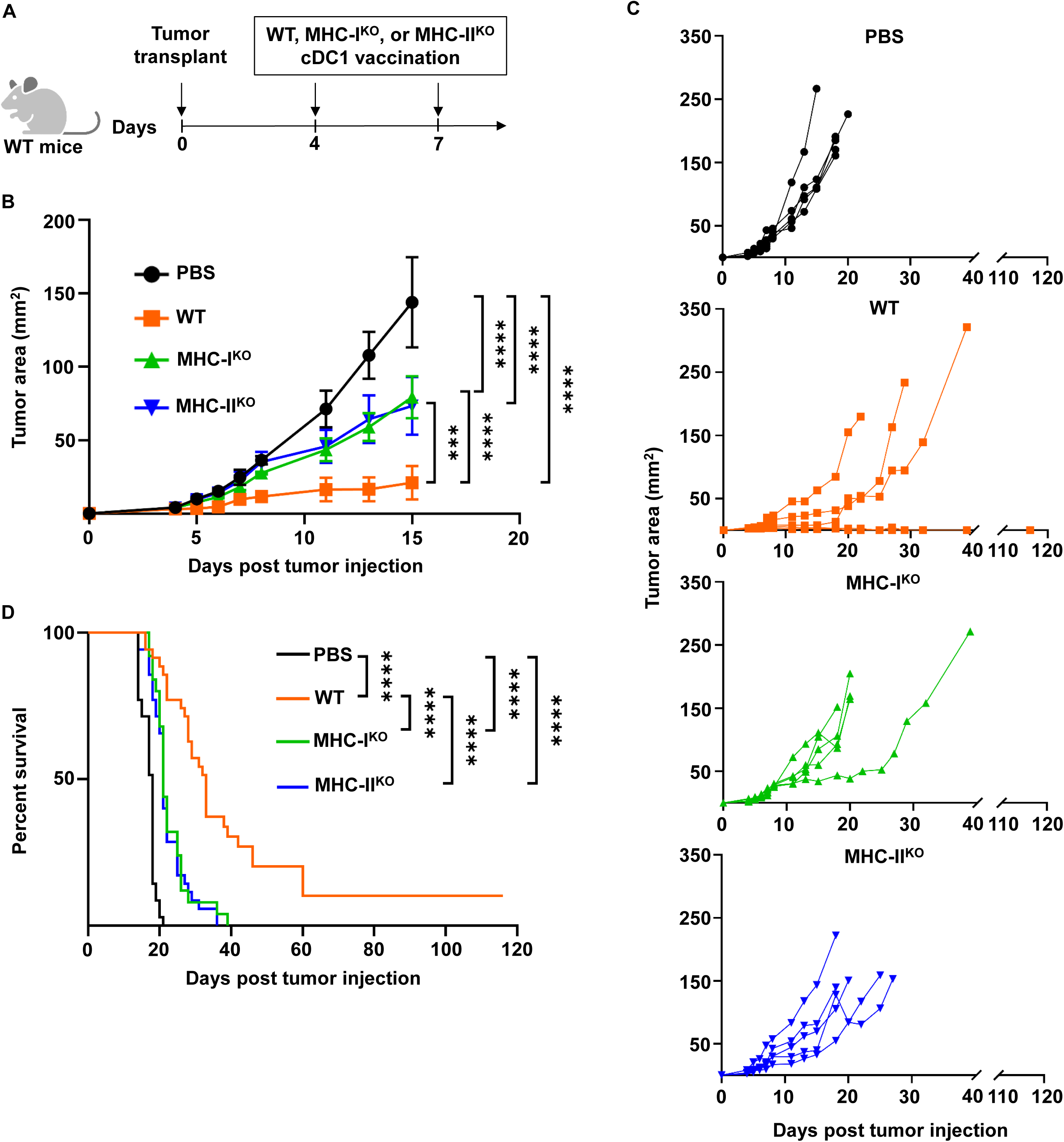
Loss of MHC-I or MHC-II antigen presentation impairs cDC1 vaccine efficacy. (**A**) Schematic of the experimental design. WT C57BL/6J mice were implanted with B16-OVA tumors on day 0 and treated intratumorally on days 4 and 7 with either PBS, WT, MHC-I^KO^, or MHC-II^KO^ cDC1 vaccines. (**B**) Representative mean tumor area (mm^2^) over time. (**C**) Individual tumor growth curves corresponding to mice in (**B**). (**B**, **C**) n = 5 mice per group; representative data from one of three independent experiments. (**D**) Cumulative mouse survival representing time to humane endpoint, defined as tumors reaching 15 mm in any direction. n = 35 (PBS), n = 35 (WT), n = 25 (MHC-I^KO^), and n = 35 (MHC-II^KO^) mice; cumulative data from at least five independent experiments. (**B**) Data shown are the mean ± S.E.M. and analyzed by two-way ANOVA. (**D**) Survival was analyzed by log-rank (Mantel-Cox) test. *** p < 0.001, ****p < 0.0001.

### Expression of MHC-I and MHC-II by the same cDC1 is critical for cDC1 vaccine efficacy

Given that loss of either MHC-I or MHC-II on vaccine-delivered cDC1s compromised tumor control, we next asked whether coexpression of both MHC molecules on the same vaccine-delivered cDC1 is important for efficacy. To test this, B16-OVA tumor-bearing mice were treated intratumorally with vaccines consisting of WT cDC1s, MHC-I^KO^ cDC1s, MHC-II^KO^ cDC1s, or a 1:1 mixture of MHC-I^KO^ cDC1s and MHC-II^KO^ cDC1s (Mix^KO^). Treatment with the Mix^KO^ cDC1 vaccine was unable to enhance tumor control relative to vaccination with MHC-I^KO^ or MHC-II^KO^ cDC1s alone and, similar to vaccines lacking MHC-I or MHC-II only, demonstrated diminished tumor control compared to WT cDC1 vaccination (**Figures 3A-3C**). Together, these results support the concept that coexpression of both MHC-I and MHC-II by the same cDC1 is required for robust tumor control following cDC1 vaccination [17, 52–54].

**Figure 3.**
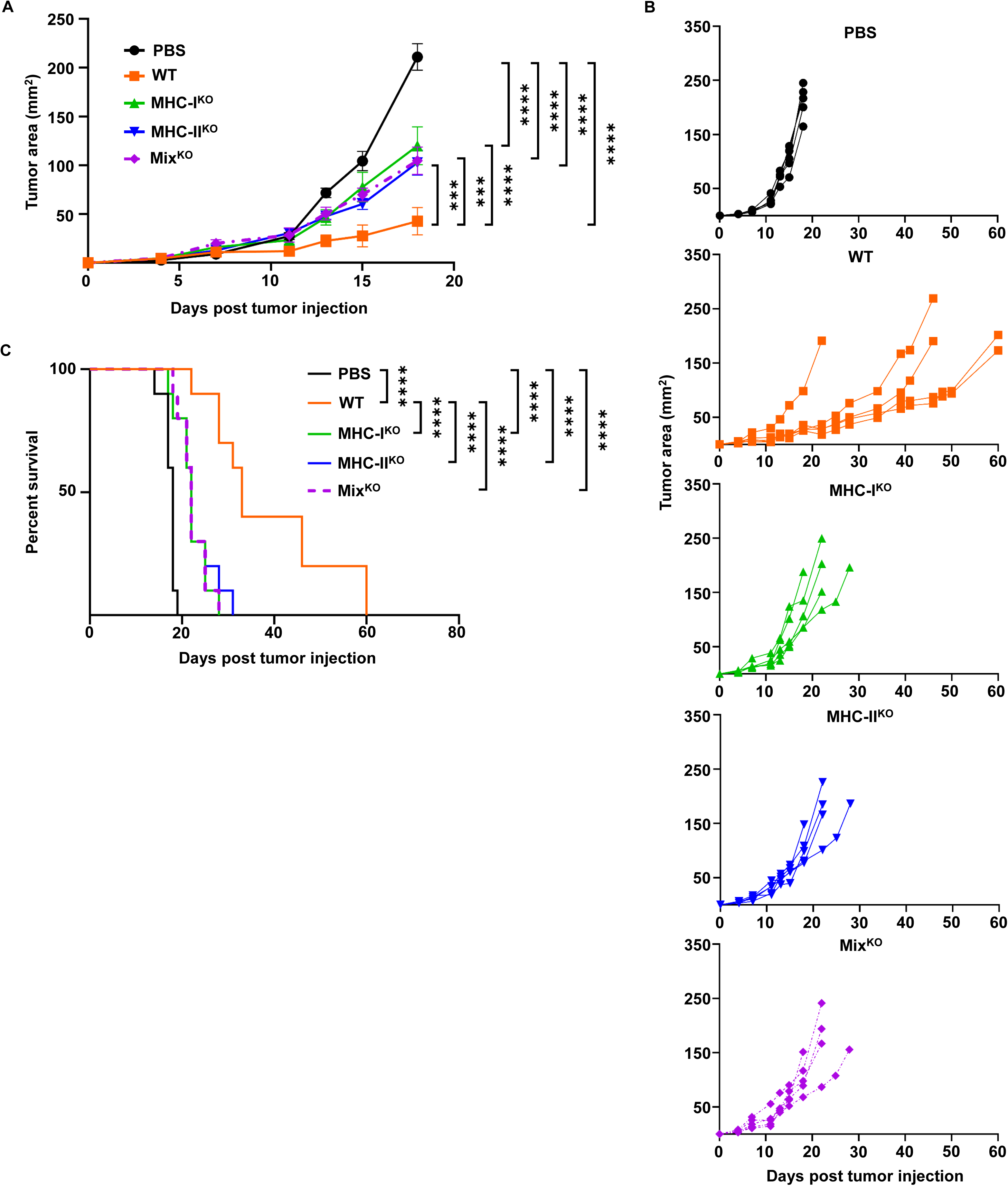
Coexpression of MHC-I and MHC-II on the same cDC1 is required for full cDC1 vaccine efficacy. WT C57BL/6J mice were implanted with B16-OVA tumors on day 0 and treated intratumorally on days 4 and 7 with either PBS, WT, MHC-I^KO^, MHC-II^KO^, or a 1:1 mixture of MHC-I^KO^ and MHC-II^KO^ (Mix^KO^) cDC1 vaccines. For Mix^KO^ vaccines, mice received 6 x 10^6^ cDC1s per injection, composed of equal numbers of MHC-I^KO^ cDC1s and MHC-II^KO^ cDC1s (3 x 10^6^ each), while individual vaccines (WT, MHC-I^KO^, or MHC-II^KO^) consisted of 3 x 10^6^ cells each. (**A**) Representative mean tumor area (mm^2^) over time. (**B**) Individual tumor growth curves corresponding to mice in (**A**). (**A, B**) n = 5 mice per group; representative data from one of two independent experiments. (**C**) Cumulative mouse survival representing time to humane endpoint, defined as tumors reaching 15 mm in any direction. n = 10 mice per group; cumulative data from two independent experiments. (**A**) Data shown are the mean ± S.E.M. and analyzed by two-way ANOVA. (**C**) Survival was analyzed by log-rank (Mantel-Cox) test. *** p < 0.001; **** p < 0.0001.

### *In vitro* CD40 ligation by agonistic antibody fails to enhance cDC1 vaccine-mediated tumor control

Since our data indicate that MHC-II is required by vaccine-delivered cDC1s for effective tumor control, we next asked whether a CD40-mediated licensing signal could enhance vaccine efficacy. We found that both WT and MHC-II^KO^ cDC1s expressed detectable levels of cell surface CD40 at steady-state and following 4 hours of poly dI:dC stimulation *in vitro* (**Figure 4A**), suggesting that both populations are capable of signaling via CD40 in the conditions used for *in vitro* activation and vaccination. Notably, CD40 expression remained detectable at 24 hours post-stimulation with poly dI:dC (**Figure S2A**), suggesting that cDC1s maintain CD40 responsiveness beyond our initial 4-hour stimulation timepoint. Furthermore, stimulation with poly dI:dC + αCD40 drove increased expression of the T cell costimulatory molecule CD70, with this effect being most evident at 16 and 24 hours post-stimulation (**Figure S2B)**, consistent with prior reports linking CD40 signaling to CD70 induction [52, 55]. We therefore tested whether exogenous activation of CD40 could bypass the need for MHC-II expression on vaccine cDC1s *in vivo* and potentially enhance WT cDC1 vaccine efficacy. B16-OVA tumor-bearing mice were vaccinated intratumorally with WT or MHC-II^KO^ cDC1s that had been treated with poly dI:dC alone or with poly dI:dC + αCD40 prior to administration (**Figure 4B**). While vaccination with WT cDC1s significantly delayed tumor progression compared to MHC-II^KO^ cDC1 vaccination or PBS treated controls, as expected, the addition of αCD40 stimulation *in vitro* did not enhance tumor control beyond that achieved with poly dI:dC-stimulated WT cDC1s (**Figures 4C-4E**). Moreover, αCD40 stimulation *in vitro* failed to improve the efficacy of MHC-II^KO^ cDC1 vaccines. Together, these results indicate that CD40 ligation provided during short-term *in vitro* activation is insufficient to enhance WT cDC1 vaccine efficacy or to compensate for the loss of MHC-II on vaccine-delivered cDC1s.

**Figure 4.**
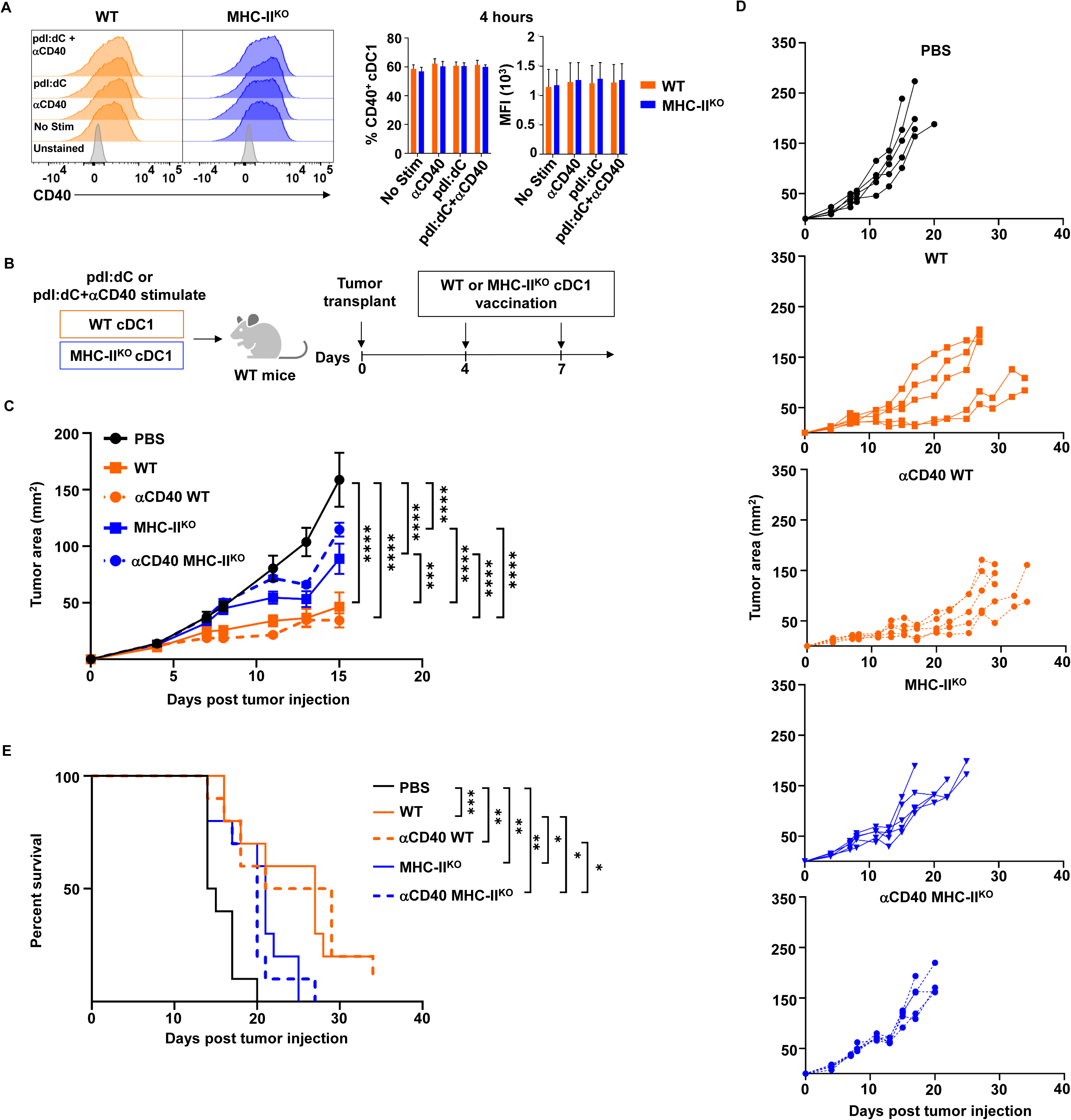
*In vitro* CD40 ligation by agonistic antibody fails to enhance cDC1 vaccine-mediated tumor control. (**A**) Representative flow cytometry histograms (left) showing CD40 surface expression on WT and MHC-II^KO^ cDC1s after 4-hour stimulation *in vitro*, with quantification of CD40^+^ frequency and CD40 MFI (right). n = 4 per group for all conditions; cumulative data from two independent experiments. (**B**) Schematic of the *in vivo* experimental design. cDC1s were stimulated *in vitro* with either poly dI:dC or poly dI:dC and αCD40 and pulsed with OVA for 4 hours, then prepared for vaccination as described in the Materials and Methods. WT mice were implanted with B16-OVA tumors on day 0 and treated intratumorally on days 4 and 7 with either PBS or cDC1 vaccines. (**C**) Representative mean tumor area (mm^2^) over time. (**D**) Individual tumor growth curves corresponding to mice in (**C**). (**C, D**) n = 5 mice per group; representative of one of two independent experiments. (**E**) Cumulative mouse survival representing time to humane endpoint, defined as tumors reaching 15 mm in any direction. n = 10 mice per group, cumulative data from two independent experiments. (**A, C**) Data shown are the mean ± S.E.M. and were analyzed by two-way ANOVA, followed by Tukey’s multiple-comparisons test applied to (**A)**. (**E**) Survival was analyzed by log-rank (Mantel-Cox) test. * p < 0.05; ** p < 0.01; *** p < 0.001; **** p < 0.0001.

### Endogenous cDC1s are required for cDC1 vaccine efficacy

Previous studies showed that cDC1 vaccination could effectively mediate tumor rejection in the absence of endogenous cDC1s, using a tumor model that shows spontaneous regression in immunocompetent hosts [56]. However, whether host cDC1s contribute to cDC1 vaccine-mediated tumor control with less immunogenic tumors remains unclear. To address this, we evaluated cDC1 vaccine efficacy against B16-OVA melanoma in *Irf8+32*^-/-^ mice (**Figure 5A**), which lack endogenous cDC1s but retain other DC lineages [34]. These studies revealed that vaccination with WT, MHC-I^KO^, or MHC-II^KO^ cDC1s resulted in reduced tumor control in *Irf8+32*^-/-^ mice compared to identically treated WT hosts (**Figure 5B; Supplementary Table 1**), indicating the presence of host cDC1s is required to achieve optimal therapeutic benefit. Notably, within *Irf8+32*^-/-^ mice, all three cDC1 vaccines still conferred a modest survival benefit relative to PBS controls, indicating that vaccine-delivered cDC1s retain partial antitumor activity even in the absence of endogenous cDC1s. Together, these data indicate that while intratumorally delivered cDC1s can partially restrain tumor progression on their own, full vaccine efficacy in poorly immunogenic tumors depends on the presence of host cDC1s, which likely provide additional immune support required for robust vaccine-elicited antitumor responses.

**Figure 5.**
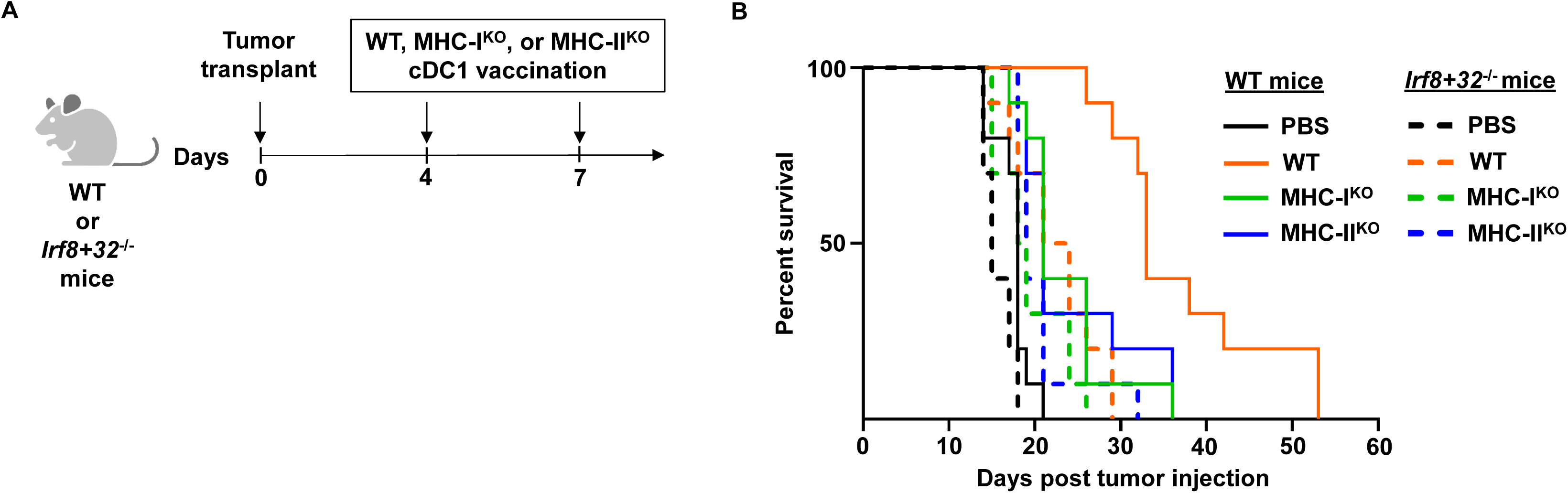
Lack of endogenous cDC1s reduces therapeutic cDC1 vaccine efficacy. (**A**) Schematic diagram of the experimental design. WT C57BL/6J or cDC1-deficient (*Irf8+32*^-/-^) mice were implanted with B16-OVA tumors on day 0 and treated intratumorally on days 4 and 7 with either PBS, WT, MHC-I^KO^, or MHC-II^KO^ cDC1 vaccines. (**B**) Cumulative mouse survival representing time to humane endpoint, defined as tumors reaching 15 mm in any direction. Survival rates of *Irf8+32*^-/-^ mice treated with the indicated cDC1 vaccines are shown as dashed lines, with identically treated WT mice shown as solid lines. Full pairwise survival comparisons are provided in Supplementary Table S1. n = 10 mice per group per genotype (WT and *Irf8+32*^-/-^), cumulative data from two independent experiments. Survival was analyzed by log-rank (Mantel-Cox) test. * p < 0.05; ** p < 0.01; **** p < 0.0001.

## Discussion

In this study, we define key antigen presentation requirements that govern the efficacy of cDC1-based cancer vaccination. By selectively disrupting MHC-I or MHC-II expression on vaccine-delivered cDC1s, we demonstrate that both antigen presentation pathways contribute to cDC1 vaccine-mediated tumor control. We further show that coexpression of MHC-I and MHC-II on the same vaccine-delivered cDC1 is required for a robust vaccine response. Notably, although host cDC1s contribute to efficacy in B16-OVA melanoma, cDC1 vaccines still elicit modest antitumor immunity in the absence of endogenous cDC1s. Collectively, these findings extend prior work establishing cDC1s as central coordinators of antitumor immunity and indicate that their capacity to simultaneously engage CD8^+^ and CD4^+^ T cells via MHC-I and MHC-II antigen presentation is functionally important in the context of therapeutic cDC1 vaccination.

Our data demonstrated that poly dI:dC stimulation of *in vitro*-derived cDC1s induced a mature cDC1 phenotype characterized by upregulation of canonical costimulatory molecules and transcriptional alterations such as induction of cytokine and chemokine gene expression. These findings raised the possibility that the costimulatory and cytokine/chemokine secretion effector functions of cDC1s might be sufficient to drive a therapeutic response following intratumoral cDC1 delivery. However, despite the similarities in these responses between WT, MHC-I^KO^, and MHC-II^KO^ cDC1s, our results show that MHC-I^KO^ and MHC-II^KO^ cDC1s have impaired ability to control tumor growth. These findings indicate that antigen presentation by cDC1s via both MHC-I and MHC-II are key determinants of vaccine efficacy.

We anticipated that vaccination with MHC-I^KO^ cDC1s would result in impaired tumor control due to the importance of cDC1s in CD8^+^ T cell priming and activation and our previous studies that linked cDC1 vaccination with increased CD8^+^ T cell responses [26]. The reduced efficacy of MHC-I^KO^ cDC1s is also aligned with the established requirement for MHC-I-mediated antigen cross-presentation by cDC1s in the priming of tumor antigen-specific CD8^+^ T cells [16, 57]. Nonetheless, MHC-I^KO^ cDC1 vaccination still provided a measurable, albeit reduced, degree of tumor control. The persistence of partial antitumor activity implies that additional mechanisms, beyond CD8^+^ T cell priming and activation, contribute to tumor control. These may include inflammatory cytokine production, chemokine-driven recruitment of immune cells, or amplification of innate antitumor responses within the TME by vaccine-delivered cDC1s, as well as the function of host cDC1s in eliciting antitumor responses. Further work to elucidate the additional factors mediating antitumor immunity may help improve strategies for effective cDC1 vaccination.

Our data also indicate that effective cDC1 vaccine responses require engagement of CD4^+^ T cells, since MHC-II-deficient cDC1s showed impaired tumor control. In addition, we found that vaccines composed of a 1:1 mixture of MHC-I^KO^ and MHC-II^KO^ cDC1s failed to restore efficacy, suggesting that interaction of both CD4^+^ and CD8^+^ T cells with the same vaccine-delivered cDC1 is necessary. These requirements are consistent with established mechanisms by which MHC-II-restricted antigen presentation by cDC1s primes CD4^+^ T cells, which subsequently deliver CD40-dependent licensing signals to cDC1s, thereby enhancing the capacity of cDC1s to prime and sustain cytotoxic CD8^+^ T cell responses [17, 52, 58]. Current evidence indicates that such CD40-dependent licensing occurs within TdLNs [17, 52, 58]. CD40 expression is enriched on migratory cDC1s in lymphoid tissues, consistent with licensing being acquired following migration from peripheral sites to LNs [17]. Additionally, CD40-dependent licensing induces key survival and functional programs in cDC1s, including upregulation of critical costimulatory ligands such as CD70, which sustain effective CD8^+^ T cell priming [52, 58]. Activated CD4^+^ T cells have also been shown to promote increases in viability, antigen presentation, and T cell activation gene programs in peripheral blood cDC1s following co-culture *in vitro* [59]. Furthermore, intratumoral immune triads composed of DCs, CD4^+^ T cells, and CD8^+^ T cells have been found to be essential for orchestrating effective antitumor responses [53, 54]. Thus, taken together, the current evidence suggests that MHC-I antigen presentation by vaccine-delivered cDC1s enables CD8^+^ T cell priming, while MHC-II antigen presentation facilitates CD4^+^ T cell engagement and cDC1 licensing signals that further stimulate CD8^+^ T cell responses, and these processes may occur in TdLNs and the TME to promote antitumor immunity.

Consistent with the above model, poly dI:dC stimulation alone of *in vitro*-derived cDC1s failed to fully induce functional features that are associated with CD40-dependent licensing. For instance, we found that αCD40 treatment was required to robustly induce expression of the costimulatory molecule CD70 on poly dI:dC-stimulated WT and MHC-II^KO^ cDC1s, in line with prior reports [52, 55], with CD70 upregulation most pronounced at relatively late time points of treatment (e.g., 24 hours). However, treatment of cDC1s with poly dI:dC and αCD40 *in vitro* (4 hours) failed to enhance the therapeutic performance of WT or MHC-II^KO^ cDC1 vaccines. Importantly, we previously established that vaccine cDC1s could be detected in tumors and TdLNs 40 hours after intratumoral delivery, indicating that a portion of the vaccine-delivered cells had prolonged survival and ability to migrate to LNs [26]. Together, these results suggest that transient CD40 stimulation, as performed in our *in vitro* stimulation conditions, is insufficient to substitute for the full spectrum of MHC-II-dependent CD4^+^ T cell help. This limitation likely reflects constraints in the timing, duration, or physiological context of CD40 signaling provided *in vitro* rather than an absolute inability of CD40 engagement to support the licensing of the cDC1 vaccine. *In vivo*, sustained interactions with CD4^+^ T cells may be required to fully induce the transcriptional, metabolic, and survival programs necessary for optimal cDC1 vaccine function [52]. CD4^+^ T cell-mediated help may also provide additional signals beyond CD40 engagement and cDC1 licensing, including locally delivered cytokines or other factors that directly regulate either cDC1s or CD8^+^ T cells, which are not fully recapitulated by short-term agonistic CD40 stimulation *in vitro*.

Our transcriptional analyses indicate that *in vitro*-derived MHC-I^KO^ and MHC-II^KO^ cDC1s retain a conserved cDC1 identity, with expression of genes involved in antigen processing, costimulation, and migration comparable to WT cDC1s. Thus, loss of MHC-I or MHC-II does not grossly alter cDC1 differentiation *in vitro*. Furthermore, when mapped onto integrated mouse and human tumor scRNA-seq datasets, *in vitro*-derived cDC1s align closely with *bona fide* intratumoral cDC1s, whereas poly dI:dC-activated cDC1s align with CCR7^+^ DCs, a DC state that is associated with maturation [49]. Moreover, expression of CCR7 on DCs is important for their migration from tumors to TdLNs, enabling the transport of tumor antigen and initiation and regulation of adaptive immune responses. Although intratumoral CCR7^+^ DCs display both maturation and immunoregulatory features, this population has been shown to contribute to activating antitumor immune responses and to associate with improved responses to immunotherapy [60]. Our data, therefore, support the notion that cDC1s generated *in vitro* recapitulate transcriptional states that naturally occur within tumors, reinforcing the relevance of cDC1-based vaccines for modeling and modulating clinically important DC states. Moreover, given that *in vitro*-derived cDC1s transcriptionally align with intratumoral cDC1s, our culture system may offer a robust approach for further dissecting physiologically relevant functions of DC states *in vivo*.

Although transcriptional profiles were largely similar among WT, MHC-I^KO^, and MHC-II^KO^cDC1s, *Btnl2* was found to be upregulated in MHC-II^KO^ cDC1s. BTNL2 has been implicated in modulating T cell costimulation [47, 48]. The increased *Btnl2* transcript expression we observed in MHC-II^KO^ cDC1s may be linked to the genomic architecture surrounding the MHC-II locus, as *Btnl2* is located adjacent to the targeted *H2-Ab1* region in the mouse genome [32, 36]. For instance, the targeted deletion may have altered regulatory elements that normally restrain *Btnl2* transcription. Despite the increased expression of *Btnl2*, we found that MHC-II^KO^ cDC1s are fully capable of inducing OT-I CD8^+^ T cell proliferation and PD-1 upregulation *in vitro*, eliciting responses comparable in extent and quality to those induced by WT cDC1s. These findings suggest that the increase in *Btnl2* does not translate into a functional defect in MHC-II^KO^ cDC1-mediated T cell activation, and that the reduced effectiveness of MHC-II^KO^ cDC1 vaccines is unlikely to result from a BTNL2-mediated impairment in CD8^+^ T activation or proliferation capacity.

Finally, we found that cDC1 vaccine efficacy in B16-OVA melanoma depended on the presence of endogenous cDC1s, as judged by impaired tumor control in *Irf8+32*^-/-^ mice [34]. These data indicate that host cDC1s provide important immune-supporting functions that aid in mounting effective antitumor responses following cDC1 vaccination. This requirement differs from other studies that have reported that cDC1 vaccines could mediate tumor rejection independently of host cDC1s in a spontaneously regressing tumor model. One possible explanation is that spontaneously regressing tumors are inherently highly immunogenic, and thus administration of exogenous cDC1s is sufficient to induce potent effector CD4^+^ and CD8^+^ T cell responses even without endogenous cDC1s. In contrast, B16-OVA melanoma tumors do not spontaneously regress and are considered poorly immunogenic. In this setting, endogenous cDC1s may be required to enhance cDC1 vaccination responses by boosting T cell priming, effector T cell expansion, and intratumoral T cell functions. Moreover, host cDC1s may become activated by sterile inflammatory factors that are released upon initial tumor killing elicited by cDC1 vaccine-induced T cell responses, thereby amplifying and sustaining T cell-mediated immunity within the tumor and TdLNs [61]. Future studies using cellular imaging or cell-tracing approaches could help further define the precise interactions and mechanisms by which vaccine-derived and endogenous cDC1s mediate antigen presentation and immune effector programming *in vivo*.

Collectively, our findings identify several key parameters relevant to effective cDC1-based cancer vaccination. First, both MHC-I and MHC-II antigen presentation must occur by the same cDC1. Second, exogenous CD40 stimulation, under the conditions tested here, is not sufficient to compensate for the absence of MHC-II expression, suggesting potential complexity in roles for CD4^+^T cell help *in vivo*. Third, endogenous cDC1s contribute substantially to vaccine performance in the poorly immunogenic B16-OVA tumor model. Together, these insights provide important design considerations for future DC-based immunotherapies, including the need to preserve MHC-I and MHC-II antigen presentation, promote CD4^+^ T cell engagement, and leverage endogenous DC function to sustain antitumor immunity.

## Lead contact

Requests for further information and resources and reagents should be directed to and will be fulfilled by the lead contact, Matthew Gubin.

## Data and code availability

## Acknowledgements

J.E.P. received educational support from Janssen Research & Development, LLC through the Janssen Scholars of Immunology Diversity Engagement Program, administered through The University of Texas MD Anderson Cancer Center. Janssen Research & Development, LLC had no role in the study design, data collection, analysis, interpretation, or decision to publish. This work was supported by the NIH grant 2R56AI109294-06A1 to S.S.W. and by CPRIT (Recruitment of First-Time Tenure-Track Faculty Members; RR 190017), an Andrew Sabin Family Foundation Fellowship, a Parker Institute for Cancer Immunotherapy (PICI) Bridge Scholar Award, a University of Texas (UT) Rising Stars Award, the University of Texas MD Anderson Cancer Center (MDACC) Support Grant (CCSG) New Faculty Award supported by the National Cancer Institute (NCI) (P30CA016672), and NCI R01CA282027 to M.M.G. M.M.G. was a Cancer Prevention and Research Institute of Texas (CPRIT) Scholar in Cancer Research and an Andrew Sabin Family Fellow during part of this study. S.M.S. is a Graduate Scholar in the CPRIT Training Program (RP210028). S.K. was a Balzan Postdoctoral Fellow supported by The International Balzan Prize Foundation. M.N.R. is a National Cancer Institute F30 Ruth L. Kirschstein Fellow (F30CA290816-01). We thank Drs. James P. Allison, Mauro DiPilato, and Kenneth Hu for advice. We also thank the MD Anderson Advanced Cytometry & Sorting Core Facility for advice and assistance which are supported by NCI Core grant P30CA016672. Graphics were created with BioRender.com.

## Authorship contributions

Conceptualization and visualization, J.E.P., S.S.W., and M.M.G.; Investigation, J.E.P., Y.Z., A.M.D., B.P., S.M.S., S.K., A.S., and M.N.R.; Formal analysis, J.E.P., T.M., L.S.; Bioinformatics analysis, T.M., L.S., J.W.; Writing-original draft, J.E.P., S.S.W., M.M.G., Writing-review and editing, J.E.P., T.M., L.S., A.M.D., B.P., S.M.S., S.K., A.S., M.N.R., J.W., S.S.W., and M.M.G. All authors have read and agreed to the published version of this manuscript.

