## Supplemental Figures for "Antigen presentation requirements for effective cDC1-based cancer immunotherapy"

Supplemental Figure 1

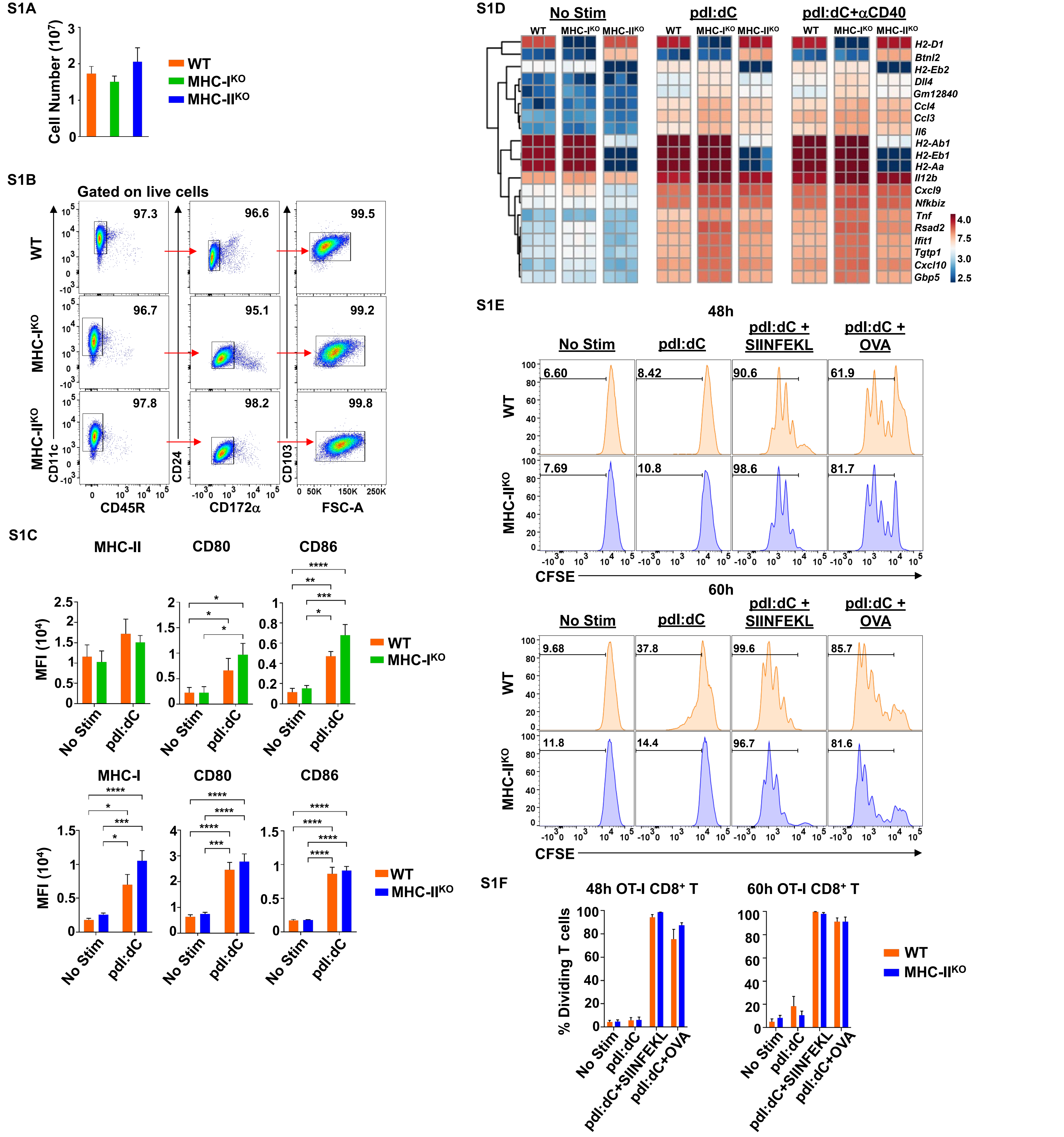

Supplemental Figure 1

S1G

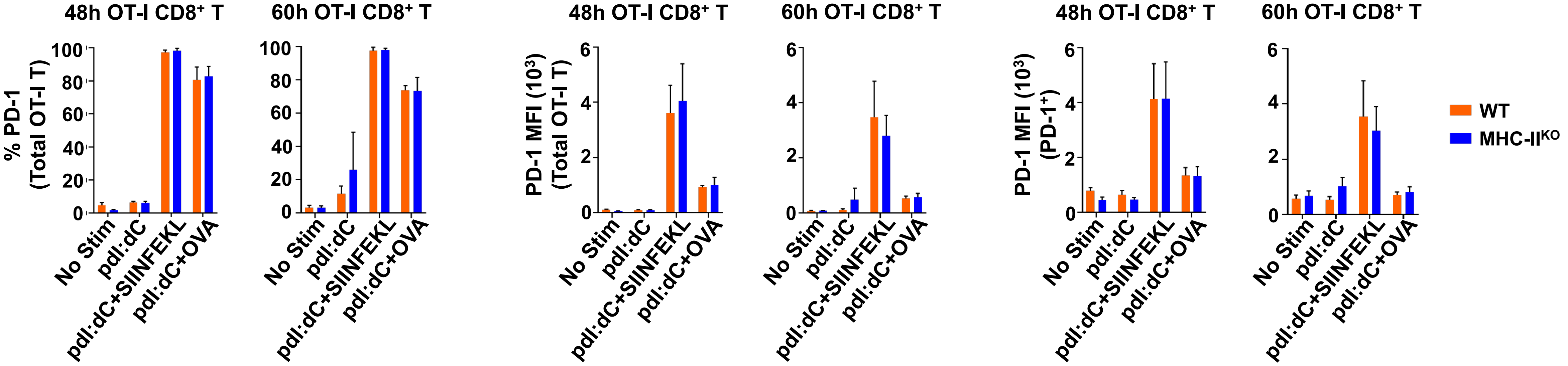


S1H

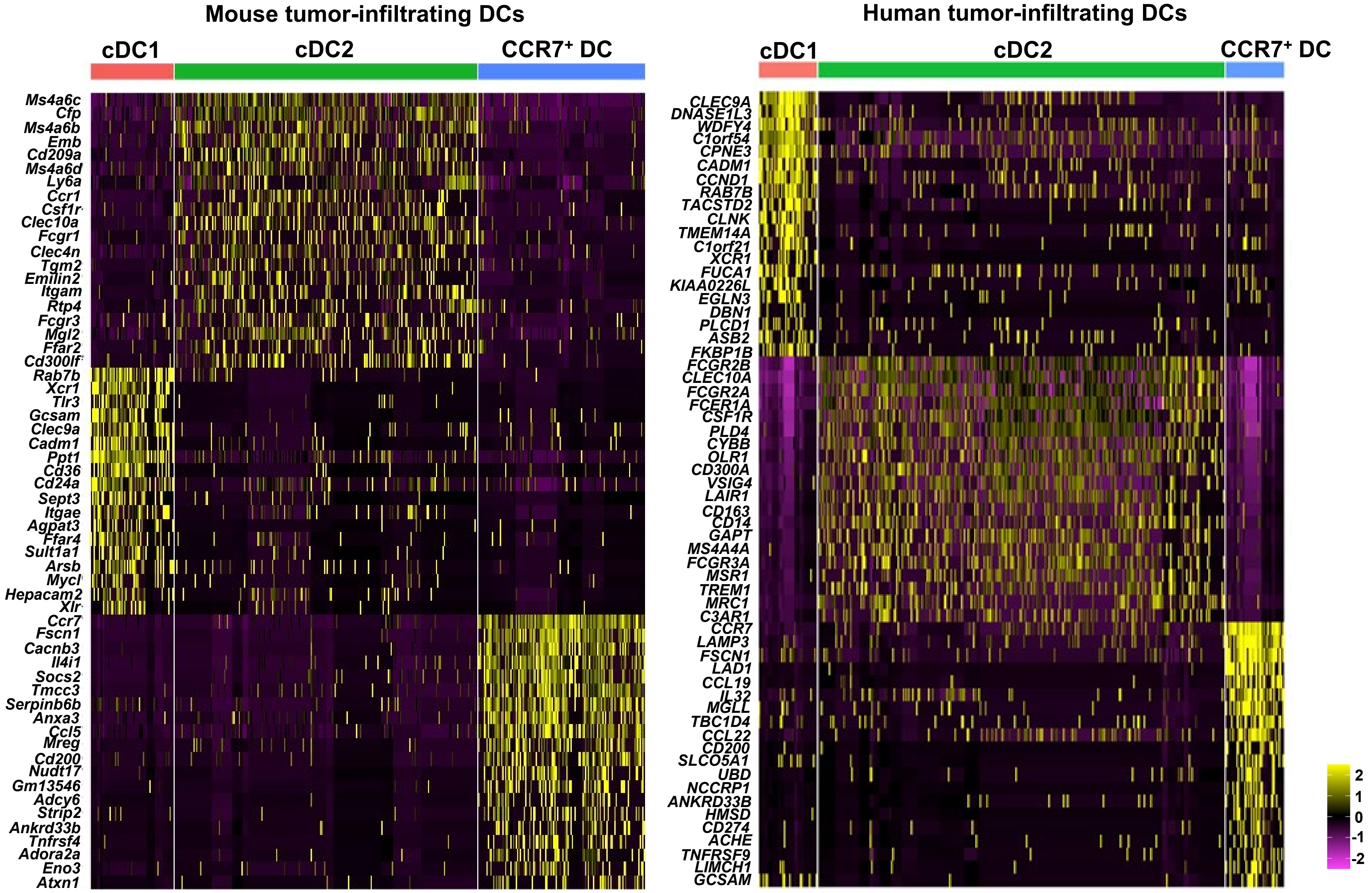

Supplemental Figure 2

S2A

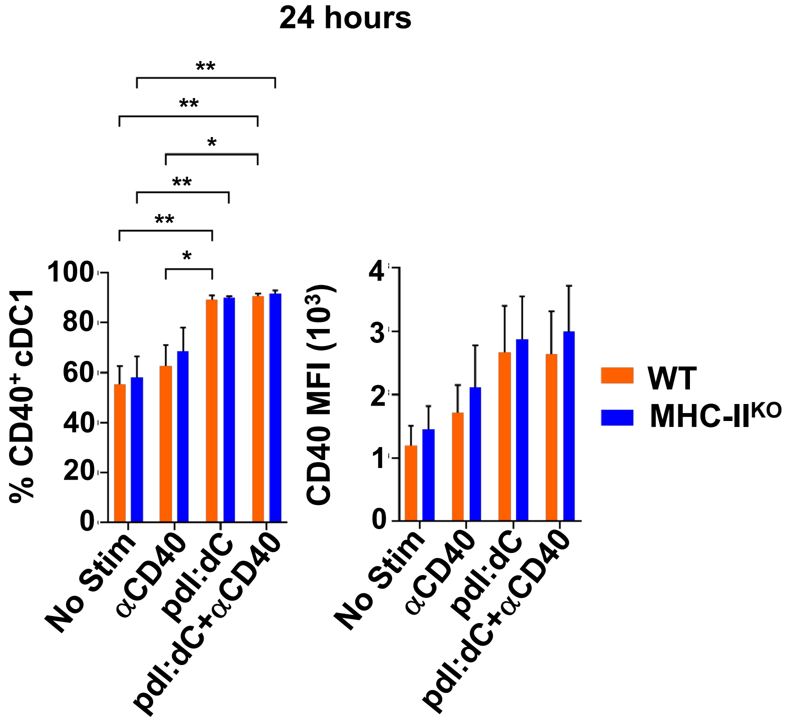


S2B

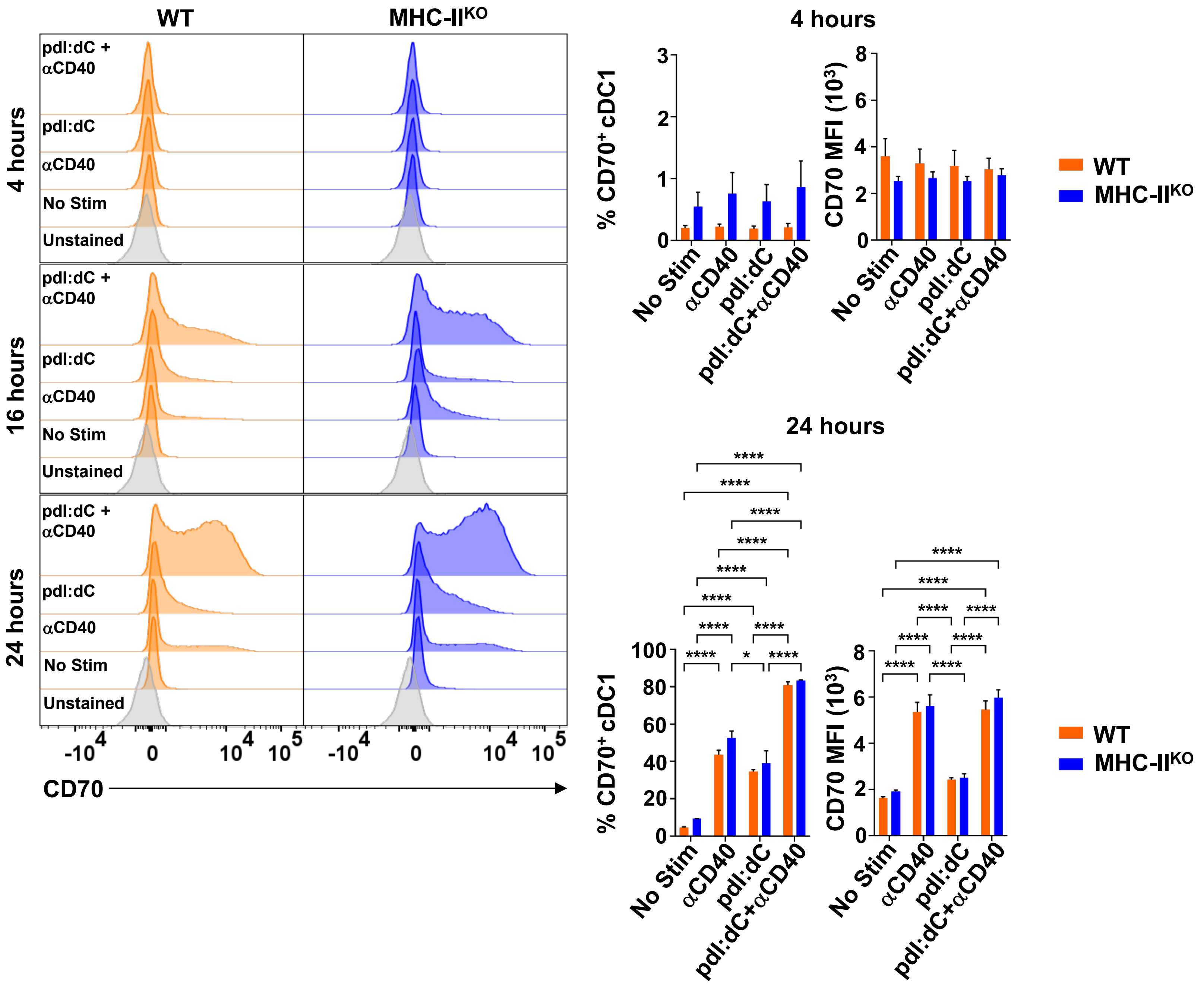

**Supplemental Table 1**

**ST1**

| **Group A<br>Mice: Vaccine** | **Group B<br>Mice: Vaccine** | **Significance** | **p value (log-rank)** |
|---|---|---|---|
| WT mice: PBS | WT mice: WT | **** | 0.000005 |
| WT mice: WT | *Irf8+32*$^{-/-}$ mice: PBS | **** | 0.000005 |
| WT mice: WT | *Irf8+32*$^{-/-}$ mice: MHC-I$^{KO}$ | **** | 0.000007 |
| WT mice: MHC-II$^{KO}$ | *Irf8+32*$^{-/-}$ mice: PBS | **** | 0.000029 |
| WT mice: MHC-I$^{KO}$ | *Irf8+32*$^{-/-}$ mice: PBS | **** | 0.000029 |
| WT mice: WT | *Irf8+32*$^{-/-}$ mice: MHC-II$^{KO}$ | **** | 0.000047 |
| *Irf8+32*$^{-/-}$ mice: PBS | *Irf8+32*$^{-/-}$ mice: MHC-II$^{KO}$ | **** | 0.000058 |
| WT mice: WT | *Irf8+32*$^{-/-}$ mice: WT | **** | 0.000059 |
| WT mice: WT | WT mice: MHC-I$^{KO}$ | *** | 0.000812 |
| *Irf8+32*$^{-/-}$ mice: PBS | *Irf8+32*$^{-/-}$ mice: WT | *** | 0.000831 |
| WT mice: PBS | WT mice: MHC-I$^{KO}$ | ** | 0.001111 |
| WT mice: PBS | WT mice: MHC-II$^{KO}$ | ** | 0.002305 |
| *Irf8+32*$^{-/-}$ mice: PBS | *Irf8+32*$^{-/-}$ mice: MHC-I$^{KO}$ | ** | 0.005424 |
| WT mice: WT | WT mice: MHC-II$^{KO}$ | ** | 0.006706 |
| WT mice: PBS | *Irf8+32*$^{-/-}$ mice: WT | ** | 0.008559 |
| WT mice: PBS | *Irf8+32*$^{-/-}$ mice: MHC-II$^{KO}$ | * | 0.021178 |
| WT mice: PBS | *Irf8+32*$^{-/-}$ mice: PBS | * | 0.024253 |
